# REMAG: recovery of eukaryotic genomes from metagenomic data using non-contrastive self-supervised learning

**DOI:** 10.64898/2026.03.05.709928

**Authors:** Daniel Gómez-Pérez, Sébastien Raguideau, Sally Warring, Robert James, Falk Hildebrand, Christopher Quince

## Abstract

Metagenome-assembled genomes (MAGs) are central to exploring microbial communities. Yet, despite the relevance of protists and fungi to diverse ecosystems, eukaryotic MAG recovery lags behind that of prokaryotes. A major bottleneck is that most state-of-the-art binning pipelines exclusively rely on prokaryotic single-copy core gene reference databases and are optimized for smaller genomes. To address this gap, we present REMAG (Recovery of Eukaryotic MAGs), a tool designed to recover high-quality eukaryotic genomes suited for long-read metagenomic data. REMAG uses a finetuned HyenaDNA genomic foundation model to efficiently filter eukaryotic contigs. It then employs a dual-encoder Siamese network trained with the non-contrastive Barlow Twins loss to learn a shared embedding space by integrating contig composition and differential coverage. Finally, high-quality bins are extracted using greedy iterative Leiden clustering optimized with eukaryotic single-copy core gene constraints. In benchmarks based on simulated mixed prokaryotic/eukaryotic communities and real datasets of varying sizes and origin, we demonstrate REMAG’s ability to recover more near-complete eukaryotic genomes than existing state-of-the-art tools, which often produce highly fragmented eukaryotic bins.

REMAG provides an automated eukaryotic binning method that scales effectively with the increasing size and sequencing depth of metagenomic datasets.

## Introduction

The popularity of metagenomic approaches for analyzing microbial communities has increased steadily in recent years, due to their potential to resolve individual genomes (Quince et al., 2017). Technological advancements that have contributed to this include long-read sequencing (Jain et al., 2016) and increases in sequencing depth from low-input material (Goodwin et al., 2016). Shotgun metagenomic sequencing data allow the building of comprehensive metagenome-assembled genome (MAG) catalogs (Singleton et al., 2026). A key step in producing these catalogs is the binning of contigs generated after metagenomic read assembly. Although state-of-the-art binners can recover near-complete bacterial genomes, this is not the case for eukaryotes, which have longer and more heterogeneous genomes (West et al., 2018). Unlike bacterial and archaeal genomes, eukaryotic genomes are typically much larger and are characterized by lower gene density, polyploidy, frequent introns and substantial repetitive content (Elliott and Gregory, 2015). Together with lower coverage, these distinct features present challenges including variable coverage patterns that can confound binners optimized to work with prokaryotic constraints.

Microbial eukaryotes have a substantial impact on the environment, driving a large part of primary production and comprising key pathogens (Parfrey et al., 2011; Falkowski et al., 2008). Despite their relevance, many are not yet sequenced due to challenges related to culturing under laboratory conditions (del Campo et al., 2014), and therefore would benefit from recovery through environmental sample collection and metagenomic analysis.

In metagenomic studies, eukaryotes have historically been neglected due to their low coverage and poor recovery. Of all the general-purpose unsupervised binners for metagenomics (CONCOCT, Alneberg et al., 2014; MetaBAT2, Kang et al., 2019; among others), none explicitly accommodate eukaryotes. Improvements in sequencing technologies and assemblers, as well as the availability of datasets enriched for eukaryotes by size filtration, such as *Tara* Oceans (Sunagawa et al., 2015), have highlighted the need for specialized eukaryotic binners. Some recently developed pipelines for short-read sequencing, such as Eukfinder (Zhao et al., 2025), are dedicated to recovering eukaryotic MAGs (eMAGs). However, they require large reference databases, which leads to slow query times and reference biases. In addition, they scale poorly with increased diversity, such as that found in newly sequenced microbial communities. Therefore, there is a need for a tool that can recover high-quality eMAGs from mixed prokaryotic/eukaryotic communities with minimal reference input.

A new generation of automated binners, including SemiBin2 (Pan et al., 2023) and COMEBin (Wang et al., 2024), is based on contrastive learning, alongside tools like VAMB (Nissen et al., 2021) that utilize deep learning to improve binning performance. Contrastive learning relies on the creation of representations that capture similarity and dissimilarity relationships among contigs (Pan et al., 2023). This is achieved by training a neural network to place data points from the same category (positive pairs) closer together in the latent space while separating those from different categories (negative pairs). In the context of binning, positive pairs correspond to versions of the same contig (halves in SemiBin2 and random subsegments in COMEBin). Negative pairs are generated randomly in both programs from different contigs assumed to be unrelated. However, these approaches are primarily optimized for prokaryotes as they rely heavily on single-copy core genes (SCGs) from archaea and bacteria for accurate clustering and do not explicitly account for the complexity of eukaryotic genomes. REMAG uses eukaryotic contig filtering prior to binning with non-contrastive Barlow Twins learning to integrate contig composition and coverage, and specifically eukaryotic marker-guided bin selection and rescue.

## Results

### Pipeline overview

REMAG implements a seven-stage integrated pipeline for eukaryotic genome recovery from metagenomic data (Figure 1). The pipeline consists of (1) eukaryotic contig filtering using a fine-tuned HyenaDNA (Nguyen et al., 2023) classifier to reduce computational load on non-target sequences and cross-domain contamination; (2) data augmentation of original contigs to generate positive pairs for training; (3) feature extraction generating compositional (tetranucleotide frequencies) and abundance (coverage) features from original and augmented contig fragments; (4) dual-encoder Siamese neural network training with the non-contrastive Barlow Twins loss to learn embeddings from both feature modalities (Figure S1); (5) k-nearest neighbor (k-NN) graph construction from learned fused embeddings; (6) greedy iterative Leiden clustering, using eukaryotic single-copy core gene (SCG) markers to select candidate bins based on estimated completeness and marker duplication; and (7) satellite rescue where fragmented bins are merged into larger core bins based on embedding centroid similarity, provided the merger does not significantly increase SCG duplication. By default, the pipeline outputs clusters with a combined length of at least 500,000 bp as bins.

**Figure 1.**
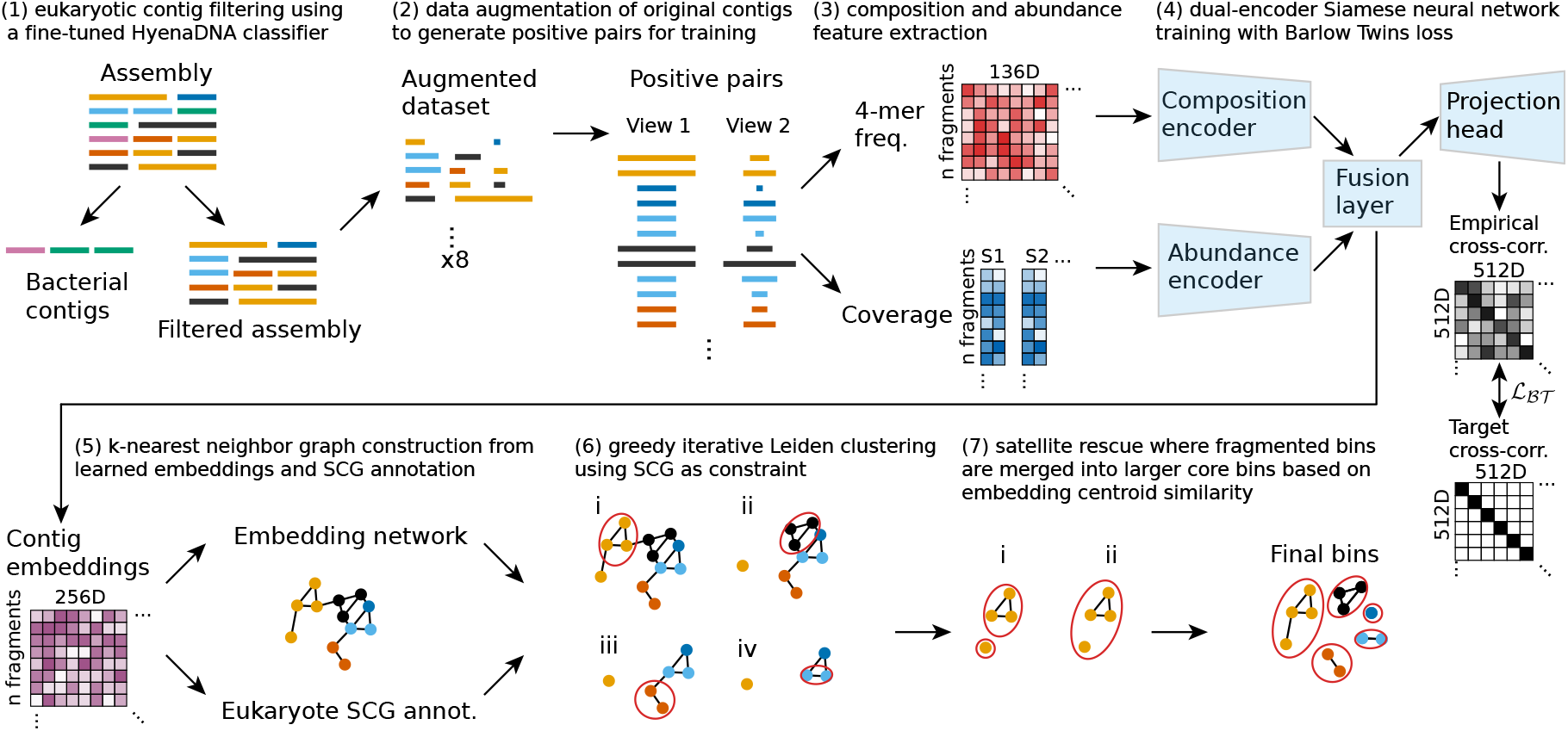
REMAG workflow. Overview of the REMAG pipeline, including (1) eukaryotic contig filtering, (2) and (3) data augmentation and feature extraction, (4) self-supervised embedding learning with Barlow Twins loss (*L_BT_*), (5) graph construction and eukaryotic single-copy core gene (SCG) annotation, (6) clustering and SCG-guided bin selection, and (7) bin rescue.

### Benchmark on synthetic datasets

We tested REMAG on mixed prokaryotic/eukaryotic communities from different biomes, including human gut, marine, plant-associated and soil for Illumina short reads and two long-read technologies, Oxford Nanopore Technologies (ONT) and PacBio. Each dataset included a variable number of low-coverage eukaryotic genomes mixed with a larger majority of prokaryotic genomes. We compared REMAG with MetaBAT2 (Kang et al., 2019), CONCOCT (Alneberg et al., 2014), SemiBin2 (Pan et al., 2023), COMEBin (Wang et al., 2024) and TaxVAMB (Kutuzova et al., 2026), using the same assembly input for each tool within the single-sample and co-assembly comparisons. We selected these tools to include established binners and recent methods that can recover eukaryotes. Figure 2 and Figure S2 use the reference-based quality tiers described in Methods.

**Figure 2.**
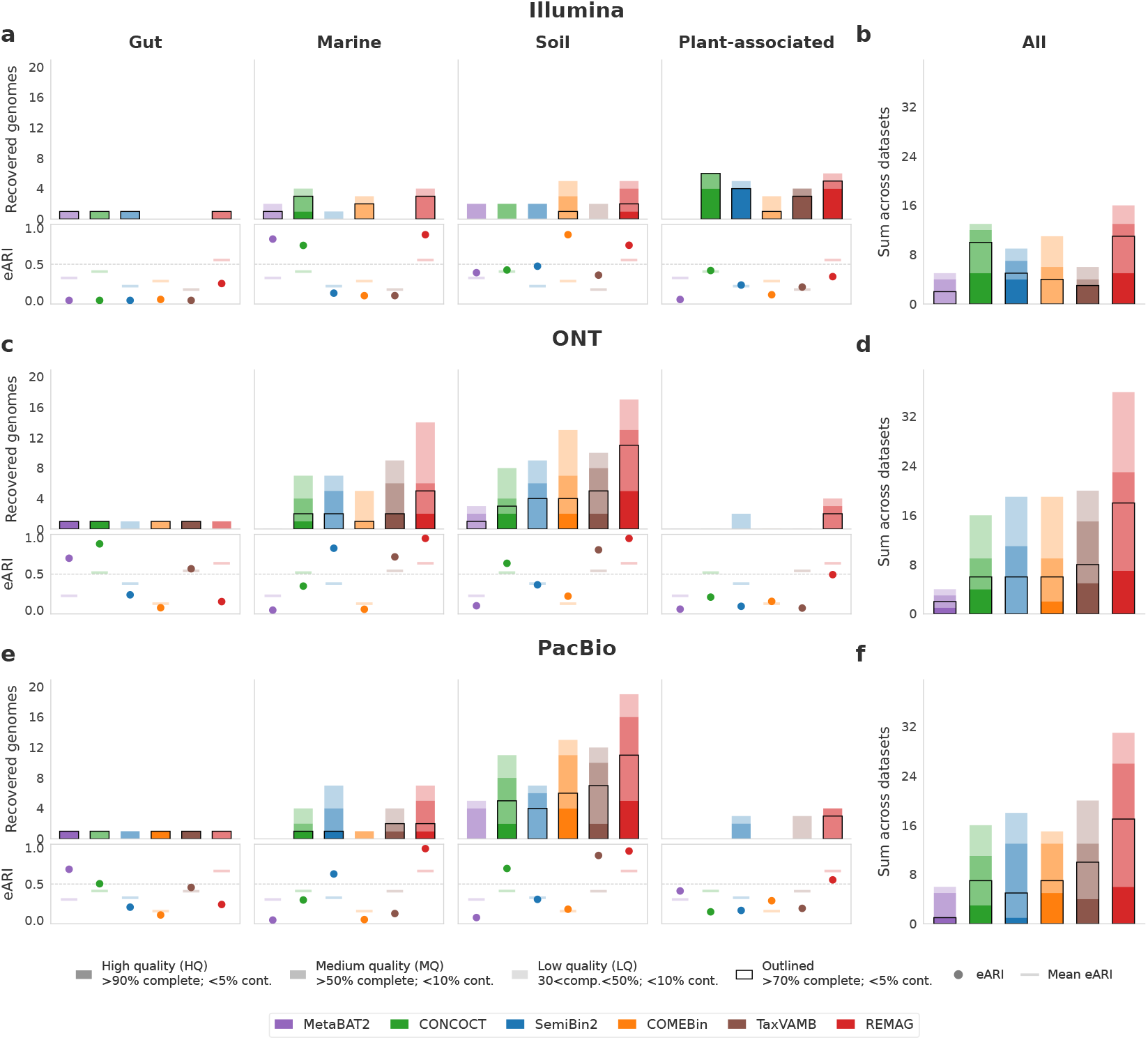
Eukaryotic genome recovery across simulated sequencing technologies. Comparison of MetaBAT2, CONCOCT, SemiBin2, COMEBin, TaxVAMB and REMAG across four biomes. Rows show Illumina (**a,b**), ONT (**c,d**) and PacBio (**e,f**). Panels **a,c,e** show unique recovered reference genomes and contig-length-weighted eukaryotic Adjusted Rand Index (eARI) for each co-assembly. Panels **b,d,f** sum recovery counts across the four datasets within each technology; these are not globally dereplicated counts. Dark, medium and light shading indicate high quality (HQ), medium quality (MQ) and low quality (LQ) tiers, respectively. Black outlines mark genomes with *>* 70% completeness and *<* 5% contamination. Points show run-level eARI, faint lines show each binner’s mean across the four datasets.

#### Eukaryotic contig filtering

As a first step, REMAG filters the assembly for eukaryotic contigs using a HyenaDNA model (Nguyen et al., 2023). This is critical to reduce the size of the input data, improving speed and reducing the potential for spurious bacterial contamination. We benchmarked our filtering approach on the coassemblies of each of the four simulated datasets by comparing performance to four other tools for eukaryotic contig classification: Tiara (Karlicki et al., 2022), 4-CAC (Pu and Shamir, 2024), EukRep (West et al., 2018) and EukFinder (Zhao et al., 2025). Contig scoring and selection were optimized for high recall to minimize loss of eukaryotic contigs. REMAG achieved the highest eukaryotic contig recall among the tested tools in all datasets except gut Illumina. Despite the higher recall at the cost of lower precision, we observed higher F1 scores for REMAG except for the gut datasets and the soil ONT dataset (Table 1). In the gut long-read datasets, eukaryotic contigs represented only approximately 0.25% of evaluated contigs, so the small proportion of prokaryotic false positives with REMAG resulted in low precision and F1 scores despite complete recall. This drop in precision is mitigated downstream, as the self-supervised learning and Leiden clustering steps effectively isolate false-positive bacterial contigs into separate bins. We compared REMAG’s learned embeddings with the input k-mer features from 4-CAC and EukRep and the learned embeddings from Tiara, and observed clearer overall separation of eukaryotic and prokaryotic contigs with REMAG (Figure S3).

**Table 1.**
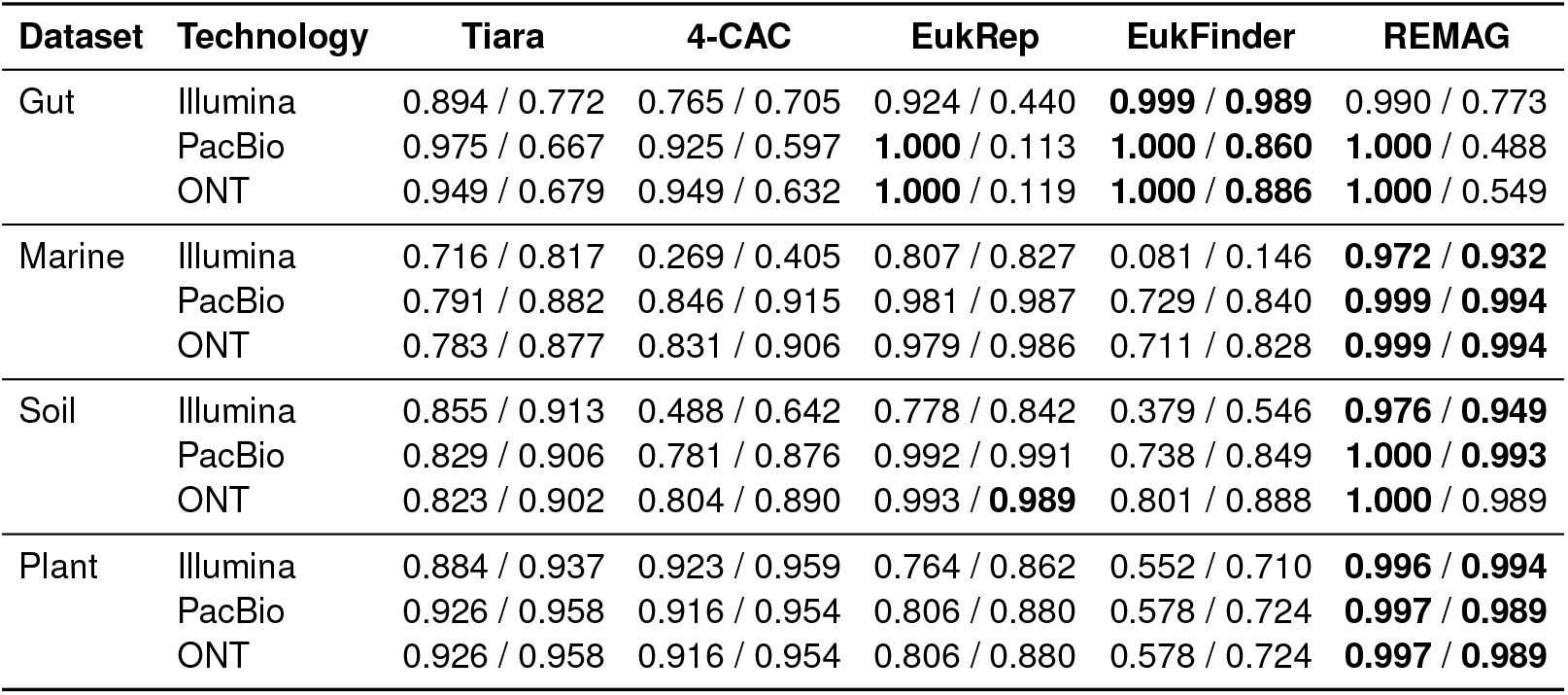
Comparison of eukaryotic contig classification on simulated co-assemblies. Each cell reports recall / F1 score, calculated from contigs at least 1000 bp long with known ground truth. Shown are results for Tiara (Karlicki et al., 2022), 4-CAC (Pu and Shamir, 2024), EukRep (West et al., 2018), EukFinder (Zhao et al., 2025) and REMAG. Within each dataset and technology, the highest recall and F1 score are bolded, based on unrounded values. ONT, Oxford Nanopore Technologies.

#### Binning on simulated single-sample assemblies

In the single-sample assemblies of the simulated datasets, all tools showed low recovery of eMAGs, with most long-read recoveries coming from the soil dataset (Figure S2). This is likely due to the low coverage of eukaryotes in these datasets, which is expected given that they were designed to reflect real microbial communities where eukaryotes are often underrepresented. Co-assembly can help recover these low-abundance eukaryotes by increasing coverage and improving assembly quality, which are critical for successful binning. Across all four datasets, REMAG recovered the most high quality (HQ) eMAGs with Illumina (2) and PacBio (7), while CONCOCT recovered the most with ONT (4 vs. 3 for REMAG). Including medium quality (MQ) eMAGs, REMAG recovered the most with Illumina (5 vs. 4 for MetaBAT2) and PacBio (17 vs. 14 for CONCOCT), whereas CONCOCT led with ONT (12 vs. 10 for REMAG).

#### Binning on short-read simulated datasets

In the short-read co-assembly simulated datasets, although optimized for long reads, we observed the best performance from REMAG. With 5 recovered HQ eMAGs, it tied with CONCOCT and recovered 2 more than TaxVAMB, while recovering 8 MQ eMAGs compared with 7 for CONCOCT across all four datasets. While CONCOCT recovered more HQ eMAGs in the marine dataset and more HQ+MQ eMAGs in the plant-associated dataset, REMAG recovered more HQ+MQ eMAGs in the soil dataset (4 vs. 2, Figure 2a,b). Of note, CONCOCT outperformed the newer tools SemiBin2, COMEBin and TaxVAMB in HQ+MQ recovery (12 vs. 7, 6 and 4, respectively).

#### Binning on long-read simulated datasets

In the longread co-assembly simulated datasets, REMAG was again the top performer, recovering the most HQ+MQ eMAGs when summed across the four datasets for each technology (23 with ONT and 26 with PacBio, compared with 15 and 13 for TaxVAMB, the next-best performer overall, Figure 2c–f). Across both long-read technologies, REMAG recovered 13 HQ eMAGs compared with 9 for TaxVAMB, with 7 HQ eMAGs recovered with ONT and 6 with PacBio. Additionally, we observed more accurate classification of contigs into eukaryotic bins on average, as measured by the eukaryotic Adjusted Rand Index (eARI). Across the eight long-read co-assemblies, REMAG had the highest mean eARI (0.67), while Tax-VAMB was second (0.47).

REMAG showed larger improvements over other tools in biomes with more diverse potentially recoverable genomes, including the marine, plant-associated and soil datasets. Across the ONT/PacBio co-assemblies, these datasets contained 7/5, 4/4 and 14/18 potentially recoverable genomes, respectively (*>*50% of reference length represented in the assembly), compared with one in each gut co-assembly. Across both long-read technologies, REMAG recovered 11, 7 and 29 HQ+MQ eMAGs in these three datasets, respectively, compared with 9, 2 and 18 for the best-performing comparator in each dataset. These results highlight the potential of REMAG for genome recovery from high-diversity environmental samples.

#### Computation time

When assessing the runtime compared to the other binners across the 12 simulated coassemblies, REMAG was the second fastest assessed binner in the GPU-assisted comparison, with a median elapsed runtime of 42.6 min, after MetaBAT2 (4.2 min, Figure S4). GPU acceleration was used for SemiBin2, COMEBin, TaxVAMB and REMAG, while MetaBAT2 and CONCOCT ran on CPUs. For a better comparison with CPU-supported binners, we reran the benchmarks using only CPUs. In this case, MetaBAT2 remained the fastest, while SemiBin2 and REMAG changed positions, with median elapsed runtimes of 65.0 min and 2.79 h, respectively (Figure S5). MetaBAT2 achieved the highest HQ and HQ+MQ eMAG recovery per runtime hour, reflecting its short runtime despite recovering only 2 HQ eMAGs (Table S1). Using the retained runtimes, REMAG ranked second for both ratios in the GPU-assisted comparison, with current recovery totals of 18 HQ and 62 HQ+MQ eMAGs.

#### Ablation testing

We tested the impact of three components in otherwise matched configurations across all 12 co-assemblies: (1) eukaryotic contig filtering; (2) the REMAG encoder, by replacing it with a simple multilayer perceptron (MLP) receiving concatenated contig composition and all coverage profiles; and (3) bin rescue. We additionally compared the encoders on 12 independently constructed five-sample co-assemblies using matched filtering and downstream settings to test encoder complexity with a low number of samples.

With filtering on, the number of recovered HQ eMAGs across all datasets and sequencing technologies increased from 12 to 18 (+50%), and HQ+MQ recovery increased from 44 to 62 (+40.9%; Figure S6). Mean eARI was approximately 0.63 in both configurations (0.629 with filtering and 0.634 without), increasing with filtering in eight of the 12 co-assemblies but decreasing in the marine Illumina and all three gut coassemblies. These results support bacterial contig removal for improving eukaryotic genome recovery overall. Filtering also reduced contamination on average among matched genomes (Figure S6c). Filtering reduced runtime in all 12 co-assemblies, with a median paired speed-up of 6.1-fold and median runtimes of 42.6 min with filtering and 291.5 min without filtering (Figure S7).

The REMAG encoder recovered 33 additional HQ+MQ eMAGs compared with the simple MLP when using all coverage profiles (62 vs. 29; +113.8%; Figure S8). Mean eARI increased from 0.42 to 0.63, improving in all 12 co-assemblies. In the five-sample co-assemblies, both encoders recovered 11 HQ+MQ eMAGs, but REMAG recovered more HQ eMAGs (6 vs. 3), with mean eARI of 0.23 compared with 0.21 for the MLP.

The rescue step also contributed to improved performance, yielding a net gain of 7 MQ eMAGs with unchanged HQ totals across the simulated datasets (62 vs. 55 HQ+MQ; +12.7%; Figure S9). Mean eARI increased from 0.55 to 0.63 (+13.9%), with increases in the marine and soil datasets (marine: 0.89 to 0.96 [+8.9%], soil: 0.69 to 0.90 [+31.6%]), although some datasets showed no change or small decreases.

### Benchmark on short-read real data

We tested REMAG on short-read datasets from the *Tara* Oceans expedition, comprising six co-assemblies from the 20–180 µm size fraction, representing surface and deep chlorophyll maximum (DCM) samples from three Pacific regions: North Pacific equatorial/tropical, North Pacific subtropical/temperate and South Pacific equatorial/tropical (Table S2). According to BUSCO, REMAG recovered 1 HQ and 25 MQ eMAGs in surface coassemblies and 26 MQ eMAGs in DCM co-assemblies, compared with 12 and 9 MQ eMAGs for CONCOCT, respectively (Figure S10). According to EukCC, REMAG recovered 3 MQ eMAGs at each depth, compared with 4 at each depth for CONCOCT and 5 and 3, respectively, for SemiBin2. While the overall recovery across all tools is limited, this likely reflects the significant divergence of these marine eukaryotic genomes from current reference databases and the inherent complexity of assembly and binning from complex marine communities using short-read sequencing.

### Benchmark on long-read real data

To demonstrate REMAG’s performance on long-read datasets, we tested REMAG on plankton sequenced with ONT and PacBio HiFi consisting of three and four samples, respectively, and assessed their quality using EukCC and BUSCO.

#### Long-read plankton datasets

REMAG recovered more HQ+MQ eMAGs than the other binners across both long-read datasets according to EukCC (10 vs. 8 for MetaBAT2, 4 for TaxVAMB, 3 for CONCOCT and 2 for SemiBin2). REMAG also recovered the most HQ+MQ eMAGs according to BUSCO (27 vs. 15 for MetaBAT2, the next-best binner). According to EukCC, REMAG recovered 6 HQ+MQ eMAGs in ONT, compared with 3 each for MetaBAT2, CONCOCT and TaxVAMB. In PacBio HiFi, REMAG recovered 4 HQ+MQ eMAGs, compared with 5 for MetaBAT2, 2 for SemiBin2 and 1 for TaxVAMB (Figure 3a,c). REMAG recovered the only HQ eMAG according to EukCC, a Bathycoccaceae bin from the ONT dataset (91.2% completeness, 2.8% contamination; Figure 3b). By contrast, a HiFi REMAG Mamiellaceae bin narrowly exceeded the MQ contamination cutoff (92.6% completeness, 10.2% contamination; Figure 3d).

**Figure 3.**
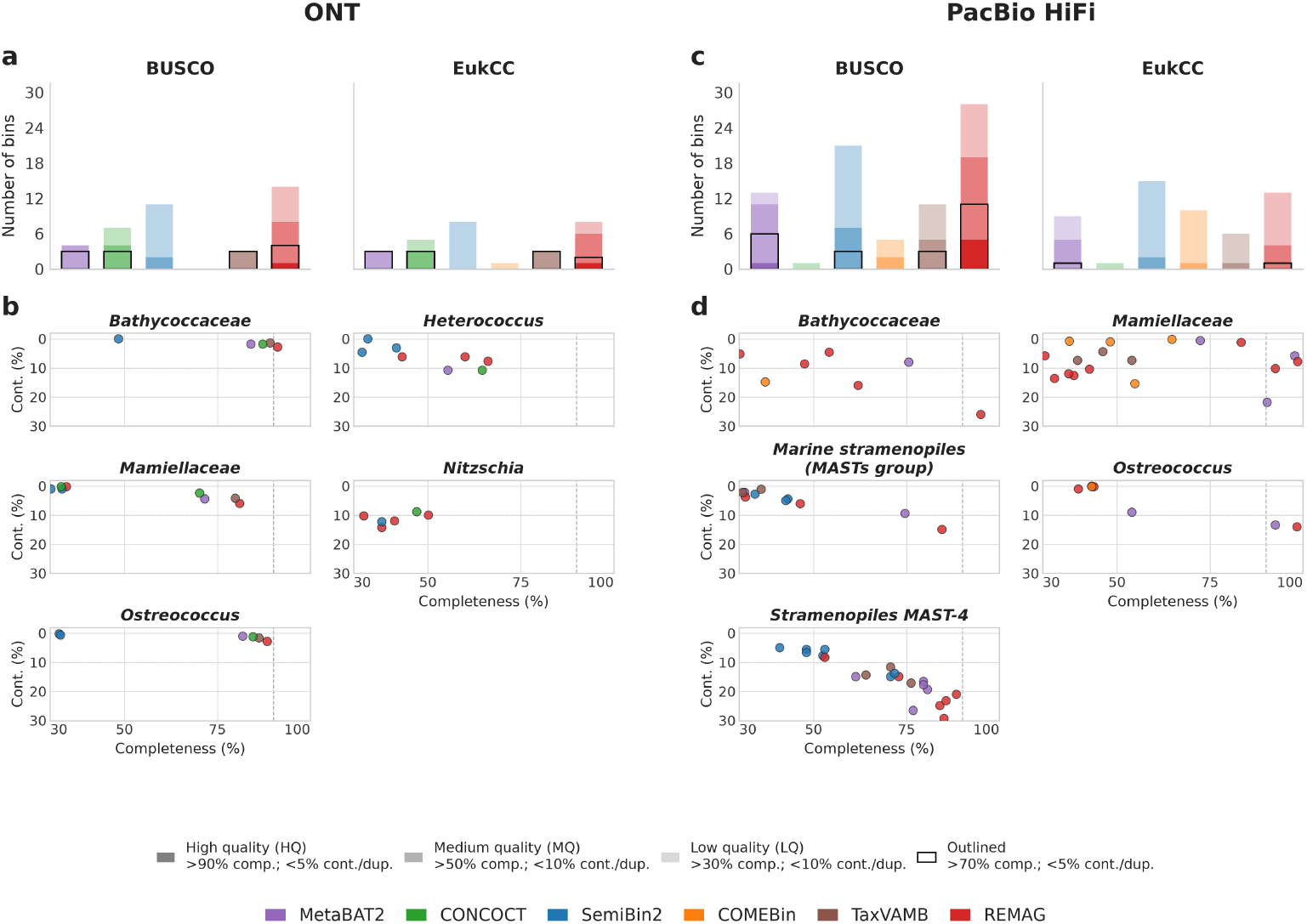
Eukaryotic MAG recovery performance on real plankton datasets. Comparison of binning tools (MetaBAT2, CONCOCT, SemiBin2, COMEBin, TaxVAMB and REMAG) on real plankton datasets sequenced with PacBio HiFi and Oxford Nanopore Technologies (ONT). **a** Shows the count of total MAGs with both BUSCO and EukCC for ONT and **b** the completeness/contamination according to EukCC for bins matched against taxonomic markers with at least one MAG classified as HQ or MQ. **c** and **d** show the results plots for the PacBio dataset. Quality tiers use EukCC contamination or BUSCO duplicated-marker percentages, as applicable.

Recovery patterns differed between sequencing technologies. According to EukCC, CONCOCT and TaxVAMB recovered more HQ+MQ eMAGs from ONT, whereas MetaBAT2, SemiBin2 and COMEBin recovered more from PacBio HiFi. REMAG performed well on both datasets, leading HQ+MQ recovery on ONT and ranking second on HiFi according to EukCC, and recovering the most HQ+MQ eMAGs on both according to BUSCO.

#### Plankton MAGs

We collated REMAG bins from both plankton datasets (ONT and PacBio HiFi) with *>*30% completeness according to EukCC. For this downstream analysis, we relaxed the contamination cutoff to *<*15% due to the presence of high-completeness bins within this contamination range (Figure 4). Dereplicating the 37 bins at 99% average nucleotide identity (ANI) resulted in 34 unique eMAGs, comprising 23 from PacBio HiFi and 11 from ONT. These eMAGs included 15 green algae, 14 marine stramenopiles (eight Ochrophyta and six MAST), one ciliate, two haptophytes and two with unresolved taxonomy.

**Figure 4.**
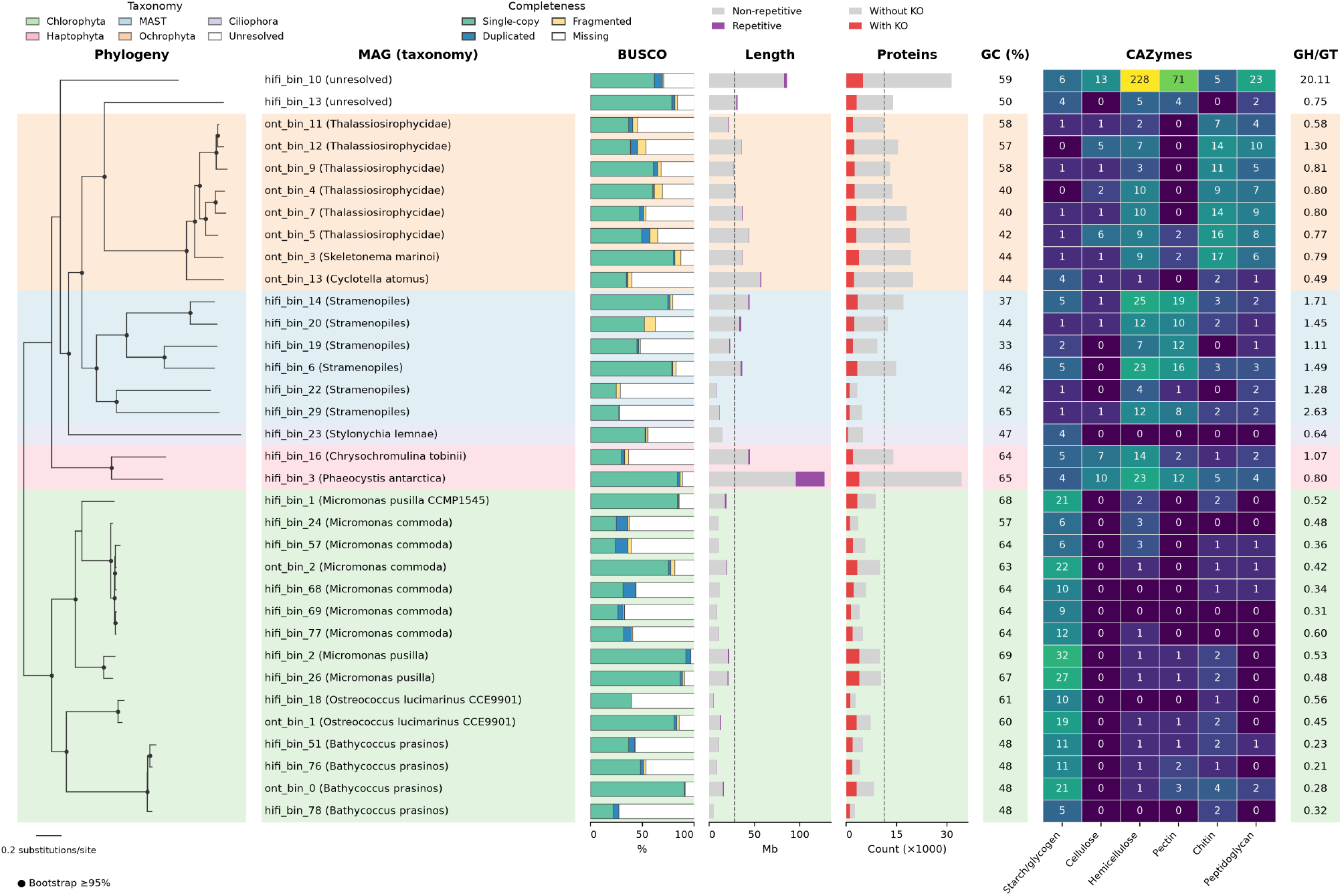
Eukaryotic MAGs recovered from plankton datasets. Phylogenetic tree of eukaryotic MAGs recovered from the plankton datasets sequenced with ONT and PacBio HiFi. Each MAG is shown with its putative taxonomic assignment in parentheses, followed by genome BUSCO completeness. Genome length, with annotated repetitive content, and the number of predicted proteins with and without KEGG Ortholog (KO) annotations are shown as stacked bar charts, with the mean shown as a dashed line. Genome GC content is shown as a percentage. The heatmap shows the number of carbohydrate-active enzymes (CAZymes) specific to different substrates. The final column gives the ratio of glycoside hydrolases to glycosyltransferases (GH/GT) in each MAG.

Core pathway completeness varied among eMAGs (Figure S11), with differences associated with recovered completeness and taxonomy. Green algae and stramenopiles also showed differences in secondary-metabolite biosynthesis modules, which may reflect their evolutionary history and niches within marine ecosystems (Amsler and Fairhead, 2005).

We also noticed differences in the distribution of carbohydrate-active enzymes (CAZymes) among the eMAGs (Figure 4). When grouped by putative substrate type, there were patterns of abundance associated with taxonomy. These included a larger number of starch/glycogen-related enzymes in green algae (mean 14.8 vs. 1.7 per eMAG in stramenopiles), which might relate to the storage of polysaccharides (Chabi et al., 2021). Stramenopiles had higher average counts of CAZymes with predicted substrates including hemicellulose, pectin, peptidoglycan and chitin, although their distribution differed between Ochrophyta and MAST. These patterns suggest potential for the degradation of complex carbohydrates, including those derived from cell walls of other organisms (Jirsová and Wideman, 2024; Christaki et al., 2017). Additionally, we found a higher glycoside hydrolase to glycosyltransferase ratio in stramenopiles compared to green algae (mean 1.14 vs. 0.41), which may reflect differences in heterotrophic potential (Seeleuthner et al., 2018).

## Discussion

The recovery of MAGs is a critical step in understanding the composition and function of microbial communities. This is particularly true of eukaryotes as, despite their relevance for the environment, they are often underrepresented in databases due to challenges with lab culturing and difficulties in generating MAGs from metagenomes. In this study, we present REMAG, a tool designed to recover eMAGs from metagenomic samples, which we have benchmarked against existing state-of-the-art binners.

REMAG uses Barlow Twins, a non-contrastive self-supervised learning objective that encourages agreement between augmented views of the same contig while reducing redundancy across embedding dimensions (Zbontar et al., 2021). Unlike contrastive-learning-based binners such as SemiBin2 and COMEBin, REMAG does not use negative pairs. This prevents noise caused by randomly sampled negative pairs that might inadvertently originate from the same genome and is particularly useful for the larger eukaryotic genomes which might be more fragmented. The exclusive use of positive links also reduces the large batch size requirements, allowing the same training volume to run faster on less memory. Furthermore, REMAG uses bidirectional cross-attention to integrate composition and coverage. A learned residual contribution and nonlinear fusion layers retain trainable integration of the two signals.

We have shown that REMAG outperforms the other tested tools in total HQ+MQ eMAG recovery from simulated co-assemblies and real long-read plankton datasets (Figures 2 and 3). The plankton eMAGs recovered by REMAG spanned a broad phylogenetic range and included both highly complete and partial genomes, with genome BUSCO completeness ranging from 24.8% to 96.8% (Figure 4). In the simulated Illumina co-assemblies, REMAG recovered slightly more HQ+MQ eMAGs than CONCOCT (13 vs. 12; Alneberg et al., 2014), with 11 and 10 eMAGs, respectively, above 70% completeness and below 5% contamination. Despite being published over a decade ago, CONCOCT remains a strong performer for eukaryotic binning in short-read datasets. This is likely because CONCOCT does not rely on single-copy markers and lacks any assumptions about genome size. REMAG’s reliance on accurate eukaryotic marker annotation for clustering likely limits its performance on short-read datasets. Shorter reads yield shorter contigs on average, potentially truncating protein-coding genes and reducing marker annotation accuracy. In the *Tara* Oceans co-assemblies, REMAG recovered the most HQ+MQ eMAGs according to BUSCO, which demonstrates its utility in real-world applications where a large diversity of eukaryotes is expected. Despite its strong performance in the simulations, TaxVAMB recovered few HQ+MQ eMAGs from the *Tara* co-assemblies. This may reflect limited representation of natural diversity in the reference databases used for taxonomic annotation (Kutuzova et al., 2026), whereas REMAG bins contigs without requiring taxonomic labels.

While REMAG has shown improvement over competing tools, the low overall recovery of eukaryotes in the real datasets demonstrates the difficulty of this problem. This is compounded by the divergence of environmental eukaryotes from reference databases, which affects both quality assessment and marker-based clustering. Future efforts should be aimed at complementing coverage and composition with further intrinsic features that provide evidence for the same-genome origin of contigs, such as assembly graphs and Hi-C contact maps.

## Conclusion

As sequencing technologies advance, the systematic study of the long and complex genomes of microbial eukaryotes from metagenomics is becoming increasingly feasible. We developed REMAG to meet the demand for microbial eukaryotic genome recovery from metagenomic data. By leveraging non-contrastive self-supervised learning and dedicated markers, it achieves superior high-quality eukaryotic MAG recovery compared to existing tools. This method is particularly valuable for exploring uncharacterized microbial diversity, as demonstrated for the plankton datasets. As metagenomic technical improvements continue and more samples become publicly available, tools like REMAG will support eukaryotic MAG recovery, complementing tools developed for prokaryotic genome recovery.

## Data and code availability

REMAG is open source and available at https://github.com/danielzmbp/remag under MIT license. An archived release (v0.6.0) is available at https://doi.org/10.5281/zenodo.23324293. The CAMI II plant-associated dataset used for benchmarking is available at https://data.cami-challenge.org/.

## Methods

### Dataset preparation

To evaluate REMAG’s performance on realistic community structures, we generated synthetic metagenomic datasets representing three distinct biomes: marine, human gut, and soil. Taxonomic profiles and abundance distributions were derived from MGnify (Mitchell et al., 2020; Table S3). The marine dataset (MGYS00001869) comprised nine samples containing 428 organisms (370 prokaryotes and 58 eukaryotes), with one, two and one unique eukaryotic genomes at least 90% represented in Illumina, ONT and PacBio co-assemblies, respectively. The human gut dataset (MGYS00002207) consisted of 19 samples with 477 organisms (464 prokaryotes and 13 eukaryotes), with zero, one and one meeting this threshold. For the soil environment (MGYS00005779), we simulated 20 samples containing 290 organisms (244 prokaryotes and 46 eukaryotes), with two, seven and seven meeting this threshold. We downloaded the corresponding genomes from GenBank and RefSeq and generated random reads using RandomReadsMG (Bushnell et al., 2025) for short-read Illumina (scaled to 35 million reads), and long-read ONT (1 million reads) and PacBio (2.5 million reads) technologies. In addition, we evaluated REMAG on the CAMI II (Meyer et al., 2022) plant-associated dataset for Illumina, ONT and PacBio, using the supplied sample-specific goldstandard assemblies for single-sample comparisons and pooled gold-standard assemblies for co-assembly comparisons, with 12, one and one genomes meeting the same threshold.

For the remaining real and simulated datasets, Illumina reads were assembled using MEGAHIT (Li et al., 2015) and mapped using BWA-MEM (Li, 2013). Long reads were assembled using metaMDBG v1.2 (Benoit et al., 2024) and mapped using minimap2 (Li, 2018).

Single-sample assemblies represent reads from one sample, while co-assemblies represent pooled reads from multiple samples within a dataset and sequencing technology. In our single-sample comparisons, each assembly was binned using its own sample’s coverage. For each co-assembly, separate coverage profiles from all contributing samples were used.

### REMAG pipeline

#### Data preprocessing

REMAG first filters for eukaryotic contigs using a single HyenaDNA tiny-16k-seqlen-d128 model (Nguyen et al., 2023), adapted to 4096-token inputs. The fine-tuning dataset comprised 31.71 million sequence fragments from 63,421 genome accessions (29,750 eukaryotic and 33,671 non-eukaryotic), split into training (85%), validation (10%) and test (5%) sets. The eukaryotic accessions comprised 13,570 fungi, 10,683 metazoans, 4,738 Viridiplantae and 759 other eukaryotes, including protists. The prokaryotic dataset contained 26,693 bacterial and 6,978 archaeal accessions. Binary classification was trained with crossentropy loss for three epochs (learning rate 2 × 10^−5^, weight decay 0.01). Using the simulated co-assembly benchmarks, we selected a mean eukaryotic probability cutoff of 0.30 and sampling of up to 16 4096 bp windows. Contigs up to 4096 bp use one padded input, while longer contigs use windows distributed evenly between both ends. Contigs meeting the minimum length (default: 1000 bp) are retained when the mean probability is ≥0.30.

#### Data augmentation

EMAG employs a random masking strategy to generate diverse training views from each contig. For contigs exceeding 50 kb, sequences are first split at the midpoint. Four masking strategies chosen at random are applied: left, right, both edges, and central masking. A maximum of eight augmented fragments are generated per contig (or contig half) by default, with a maximum 25% overlap constraint between fragments to ensure diversity and reduce training on redundant features. Contigs and training fragments must meet the minimum length threshold (default: 1000 bp). The process randomly creates training pairs where augmented views from the same contig and the original contig serve as positive examples (default: maximum 5,000,000 pairs). These positive pairs include all combinations of the original contig and augmented fragments.

#### Feature extraction

For each original contig and fragment, two feature modalities are computed: composition and abundance. Compositional features consist of canonical tetranucleotide (4-mer) frequencies (136 dimensions) calculated using reverse-complement aware counting, where each 4-mer and its reverse complement map to the same feature dimension (Dick et al., 2009). The 136 dimensions represent the 256 possible 4-mers reduced by palindromes and reverse complement equivalence (Nissen et al., 2021). Frequencies are normalized by fragment length with pseudocount addition (1e-5) to avoid zero-division.

REMAG accepts BAM/CRAM alignment files and precomputed contig-level or interval coverage files. For alignment data, per-base coverage is computed using pysam, a wrapper for SAMtools (Li et al., 2009), then averaged across fragment coordinates. Both mean coverage and standard deviation are computed per contig or fragment per sample. These raw coverage values are first normalized by the total mapped reads per sample (converting to reads per million) to account for differing sequencing depths. All normalized coverage values then undergo log1p transformation followed by global MinMax scaling to [0,1] range, preserving coabundance patterns across samples.

#### Modal encoders

The network architecture consists of a dual-encoder Siamese network with separate MLPs for 4-mer frequencies (composition) and coverage (abundance) features. The composition encoder uses two fully connected layers (136 → 128 → 64) with batch normalization, LeakyReLU activation, and 5% dropout. The abundance encoder uses the same structure (*p* → 64 → 32), where *p* is the number of coverage features.

#### Fusion layer

A fusion layer projects both modalities to a shared embedding space (default: 256 dimensions) using a bidirectional cross-attention mechanism (Vaswani et al., 2017; Chen et al., 2021). The fusion architecture employs multi-head attention modules (four heads, 10% dropout) enabling both 4-mer and coverage features to attend to each other, capturing complementary compositional and abundance information.

For each modality, the projected features *h* are averaged with their cross-attended context to give *z*. A learned residual weight *ρ* retains the original signal: *z^′^* = BN ((1 − *ρ*)*z* + *ρh*), where BN denotes batch normalization. The weight is initialized to 0.5 and constrained to 0.1–0.9.

To model patterns at different granularities, we apply a multi-scale feature module to both gated modality vectors. The module uses three parallel transformations: a bottleneck path (*E* → ⌊*E/*2⌋ → *E*), a same-scale linear path (*E* → *E*), and an expansion path (*E* → 2*E* → *E*), where *E* is the embedding dimension. At each scale, transformed k-mer and coverage vectors are concatenated, and all scale-specific outputs are concatenated into a single multi-scale representation. This representation is then compressed and passed through a non-linear interaction block, followed by adaptive dropout. Finally, the interaction output is concatenated with the gated modality vectors and the cross-attention alignment signal before the final fusion MLP. For datasets lacking coverage data, 4-mer features bypass the crossattention stage and are processed through a simple projection layer.

#### Projection head and training

A projection head (256 → 128 → 512 dimensions) with batch normalization and ReLU activation transforms embeddings for loss calculation during training. The network is trained exclusively on positive pairs using the non-contrastive Barlow Twins loss (Zbontar et al., 2021) (default: *λ* = 3 × 10^−3^), which encourages invariance across augmented views while reducing redundancy across embedding dimensions through cross-correlation matrix regularization. The loss function is defined as:

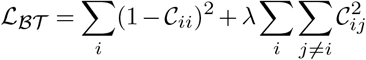

where *C* ∈ ℝ*^d×d^* is the cross-correlation matrix, computed across the batch dimension from the batch-normalized embeddings of the two identical networks, and *d* is the projection head output dimensionality. The first term minimizes the difference between diagonal elements and 1 (invariance objective), while the second term minimizes off-diagonal elements (redundancy reduction objective).

Optimization employs AdamW (Loshchilov and Hutter, 2017) (*β*_1_=0.9, *β*_2_=0.95, weight decay=0.05) with gradient clipping (max norm=1.0). Learning rate scheduling includes a 10-epoch linear warmup from 10% to full learning rate, followed by cosine annealing to 1% of peak rate. The benchmarks used a base learning rate of 5 × 10^−4^, linearly scaled by (*batch*_*size/*256) × 0.2. Training used the default batch size of 2048 for up to 100 epochs with early stopping monitoring training loss (patience=20 epochs).

For the encoder ablation, composition and coverage features were concatenated and passed through two linear layers (136 + *p* → 64 → 256, where *p* is the number of coverage features) with ReLU between them and no encoder normalization, dropout, attention or fusion modules.

#### Graph construction and marker annotation

A k-NN graph is constructed from L2-normalized embeddings resulting from the fusion layer (default: 256 dimensions) using cosine similarity. REMAG uses a minimum contig length for graph construction. For multi-sample inputs, the default graph threshold is 3000 bp. Shorter contigs remain available for rescue after clustering. The default parameters include k=15 neighbors per node and a similarity threshold of 0.1 to prune weak edges. Graph construction is parallelized using sklearn’s NearestNeighbors (Pedregosa et al., 2011) with brute-force search. The resulting undirected graph uses cosine similarity as edge weights. All contigs are annotated by mapping the 133 families of eukaryotic SCGs from ORTHODB version 12 (Kuznetsov et al., 2023) using Miniprot (Li, 2023).

#### Greedy iterative clustering

REMAG employs a greedy iterative clustering strategy to identify high-quality bins while avoiding the limitations of a single global resolution parameter. The algorithm iteratively extracts bins from the k-NN graph using the following process. (1) The current graph (initially containing all contigs meeting the graph length threshold) is clustered using the Leiden algorithm (Traag et al., 2019) with cosinesimilarity edge weights across a range of resolutions (default: 0.1, 0.5, 1.0, 2.0, 5.0). (2) For every cluster generated at each resolution, a quality score is calculated based on SCGs. The F1 score is the harmonic mean of completeness (*C* = *N*_unique_*/*133) and precision (*P* = 1 − *K*), where *K* is contamination defined as the fraction of total marker genes found (*G*) that are redundant 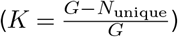:

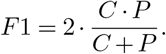

(3) Before selection, candidate clusters are filtered using a strict duplication threshold; clusters where the number of duplicated SCG families exceeds 10% of the total 133 marker families 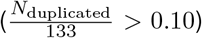 are rejected. The valid cluster with the highest F1 score across all tested resolutions is then selected as a bin, and the contigs belonging to this bin are removed from the graph. (4) The process repeats with the remaining nodes until no further valid bins can be formed.

#### Satellite rescue

After clustering, REMAG merges bins that may have been over-fragmented. This step aims to merge smaller “satellite” bins into larger “core” bins based on their embedding similarity, subject to contamination constraints. Each bin’s centroid is the mean of its high-dimensional contig embeddings, weighted by sequence length. Each bin is considered for merging, from smallest to largest. The algorithm searches for a target bin among larger or equally sized bins that shares the highest cosine similarity with the candidate’s centroid, provided the similarity is at least a defined threshold (default: 0.70). A merge requires at least one edge in the k-NN graph connecting the current source and target bins. The SCG duplication rate 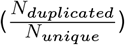 is calculated for the merged bin. By default, merged bins must have ≤5% SCG duplication. At high cosine similarity (≥ 0.95), a target already above this limit accepts a whole-bin merge only if duplication does not increase. Unassigned graph contigs and shorter contigs are recruited at the same default similarity threshold.

### Recovery quality assessment

Following established completeness and contamination thresholds (Bowers et al., 2017), high quality (HQ) MAGs were defined as having *>*90% completeness and *<*5% contamination. Medium quality (MQ) MAGs were defined as having *>*50% completeness and *<*10% contamination, excluding HQ bins. Bins with *>*30% and *<*50% completeness and *<*10% contamination were classified as low quality (LQ). For datasets with ground truth, we mapped the binned contigs back to the reference genomes using minimap2 (Li, 2018) and generated a contig-to-genome mapping file for each dataset. For real datasets, we used BUSCO (Manni et al., 2021) with the eukaryota_odb12.2 database and Eu-kCC (Saary et al., 2020) for quality assessment.

For the simulated datasets, recovery was scored using the best bin for each reference genome. Completeness was the union of covered reference bases divided by full reference-genome length. Contamination was the fraction of non-target bin bases. For the single-sample simulations, each reference genome was counted once at its best quality across samples within a dataset, sequencing technology and binner.

Clustering accuracy was quantified using the Adjusted Rand Index (ARI) computed from ground-truth contig-to-genome assignments and predicted bin labels. We report a eukaryote ARI (eARI) weighted by contig length. This eARI includes only bins that contain at least one eukaryotic contig, plus unbinned eukaryotic contigs, to penalize bins that mix eukaryotic and non-eukaryotic content.

### Plankton MAG downstream analysis

For downstream characterization of plankton MAGs, we generated a phylogeny from single-copy marker proteins annotated with BUSCO (Manni et al., 2021) and the eukaryota_odb10 lineage present in more than 50% of MAGs, due to the larger pool of markers compared to ODB12. Marker protein sequences were aligned with MAFFT (Katoh and Standley, 2013), and a phylogenetic tree was inferred with IQ-TREE v3 (Wong et al., 2025). Gene calling was performed with BRAKER3 (Gabriel et al., 2024) on the repeat-masked (Smit et al., 2013–2015) contigs, retaining one longest predicted protein per gene. Taxonomy assignment was performed using MMseqs2 (Steinegger and Söding, 2017) against UniRef90 (The UniProt Consortium, 2023). Functional annotation was performed by assigning KEGG Orthologs (KOs) with KofamScan using the August 2026 profile HMMs and adaptive score thresholds (Aramaki et al., 2020). KEGG module completeness was calculated with the EBI KEGG module graph tool (Richardson et al., 2023). CAZymes were annotated using dbCAN (Zheng et al., 2023) with the 2024 database.

### Computational resources

All analyses were executed on a single node equipped with 16 CPU cores, 128 GB RAM and an NVIDIA A100 GPU with 80 GB of memory.

## Acknowledgements

This work was supported by the BBSRC (UKRI) through the Earlham Institute Strategic Programme Grant BBX011089/1 (WP1: BBS/E/ER/230002A) and Core Capability Grant BB/CCG2220/1. We thank the Earlham Institute Research Computing Group for providing High Performance Computing resources.

## Supplementary Information

**Figure S1.**
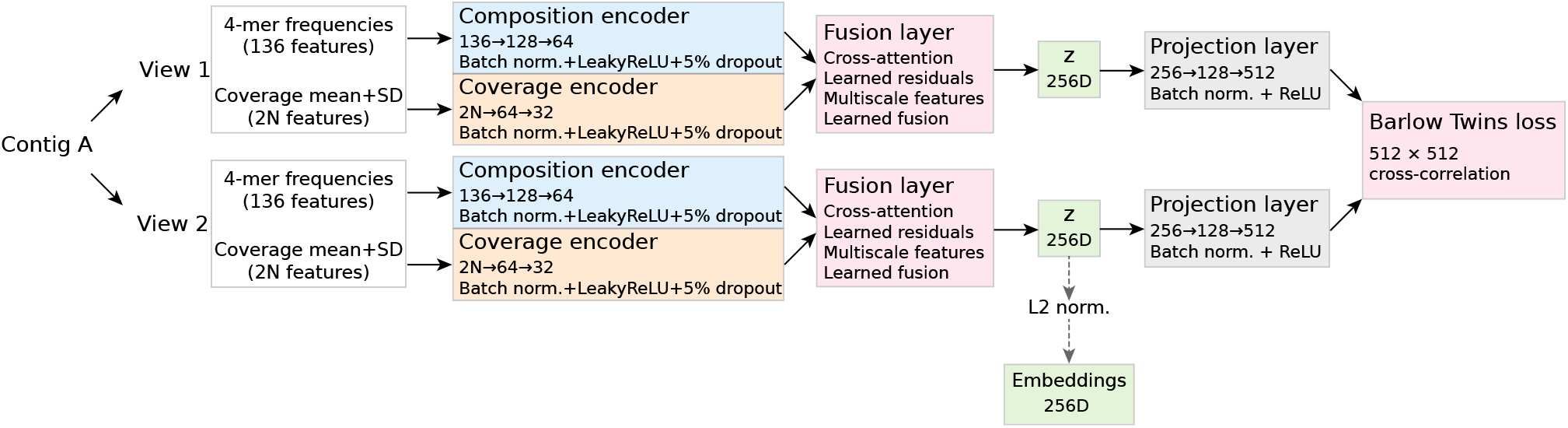
REMAG model architecture. Two views of the same contig pass through a shared network combining composition and coverage features. The fused embeddings are projected for training with Barlow Twins loss and L2-normalized for clustering. Dashed arrows show embedding extraction from original contigs after training. N denotes the number of coverage profiles.

**Figure S2.**
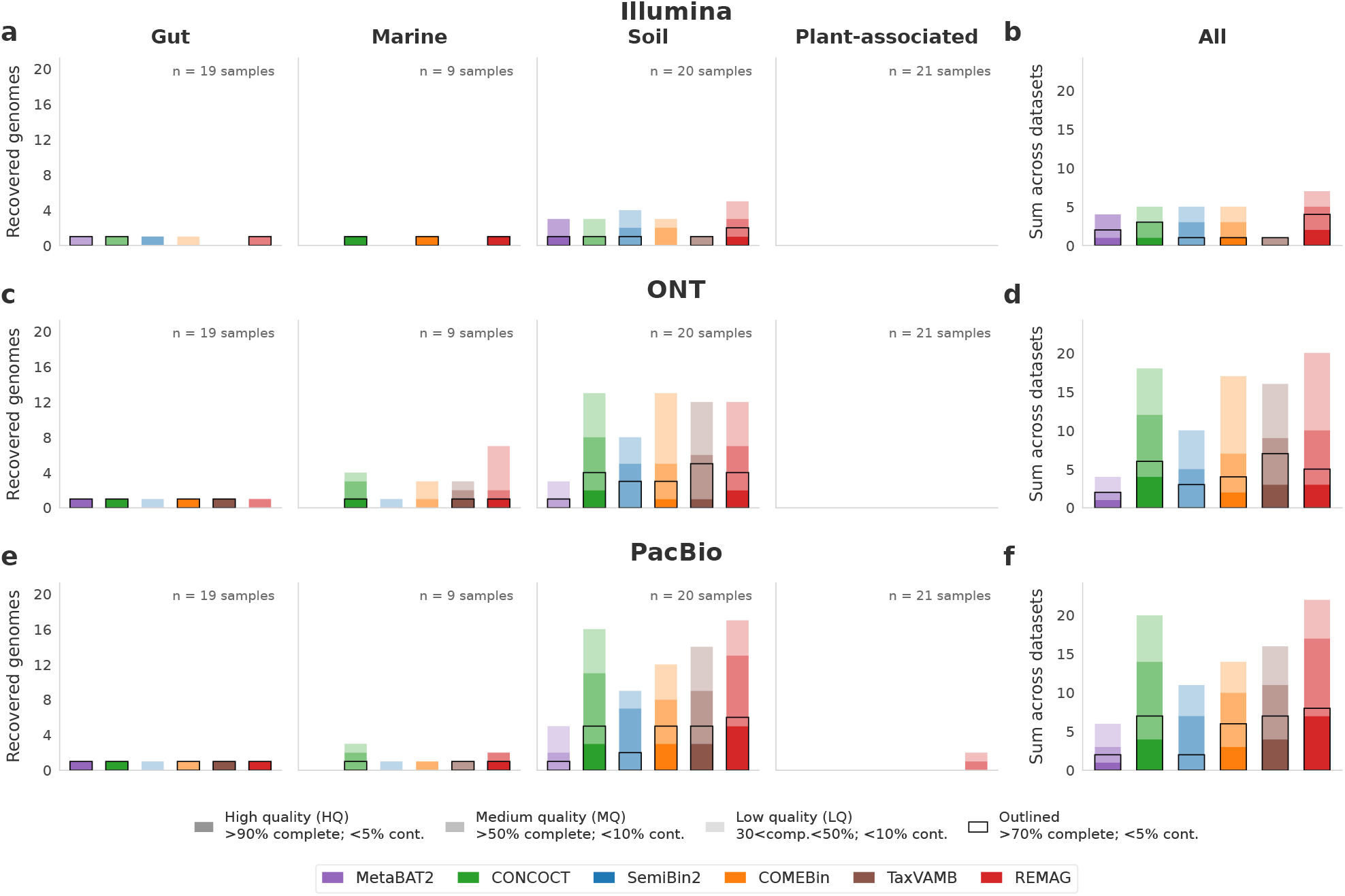
Unique eukaryotic genome recovery from single-sample assemblies. Comparison of MetaBAT2, CONCOCT, SemiBin2, COMEBin, TaxVAMB and REMAG across 207 simulated assemblies from different biomes. Rows show Illumina (**a,b**), ONT (**c,d**) and PacBio (**e,f**). Panels **a,c,e** show recovery by dataset, with the number of samples indicated. Each reference genome is counted once at its best quality tier across samples within a dataset, sequencing technology and method. Panels **b,d,f** sum these dataset-level counts.

**Figure S3.**
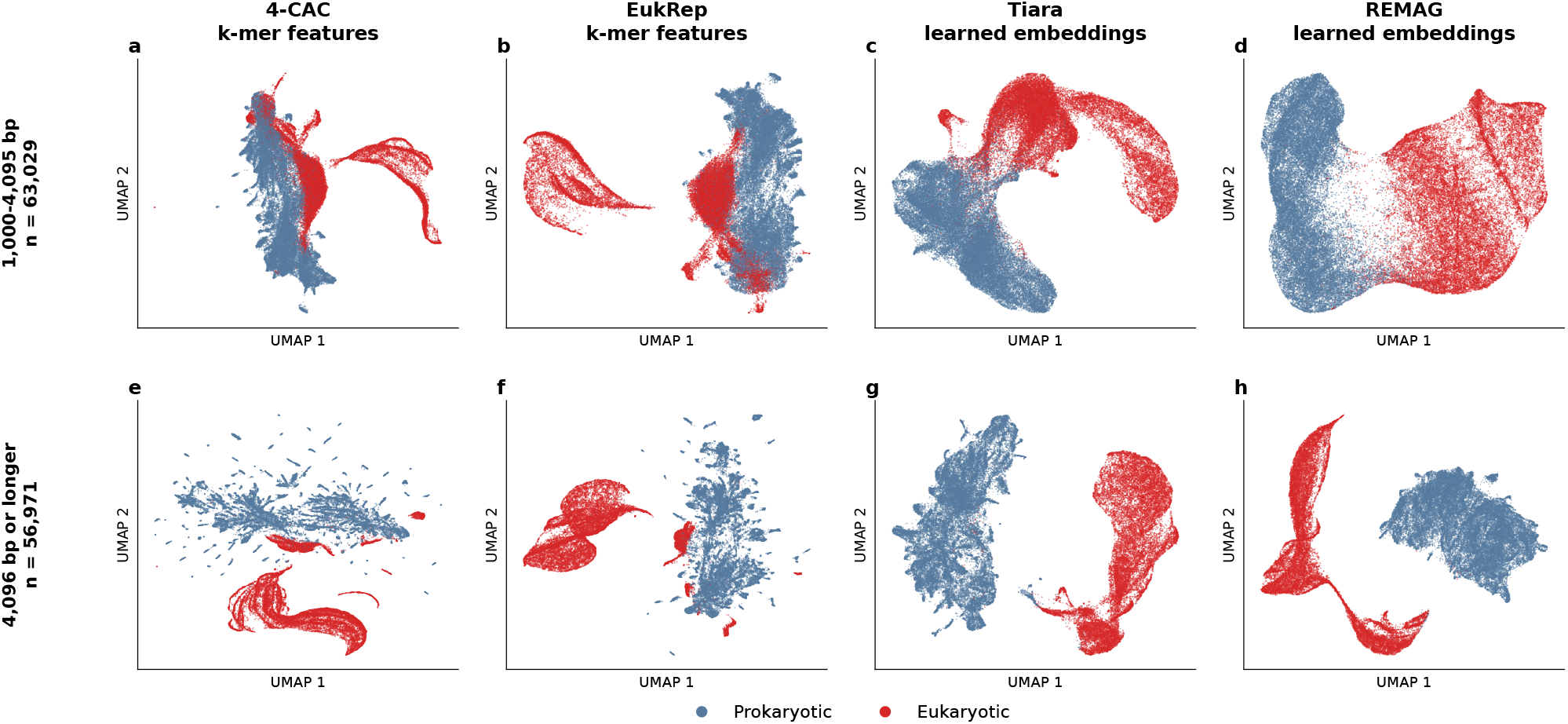
Contig representations for eukaryotic classification. UMAP projections of 10,000 randomly selected contigs from each of the 12 simulated co-assemblies, coloured by eukaryotic or prokaryotic origin. The same contigs are shown for all tools. Rows separate contigs of 1,000–4,095 bp and *≥*4,096 bp. 4-CAC and EukRep show input k-mer features, while Tiara and REMAG show learned embeddings.

**Figure S4.**
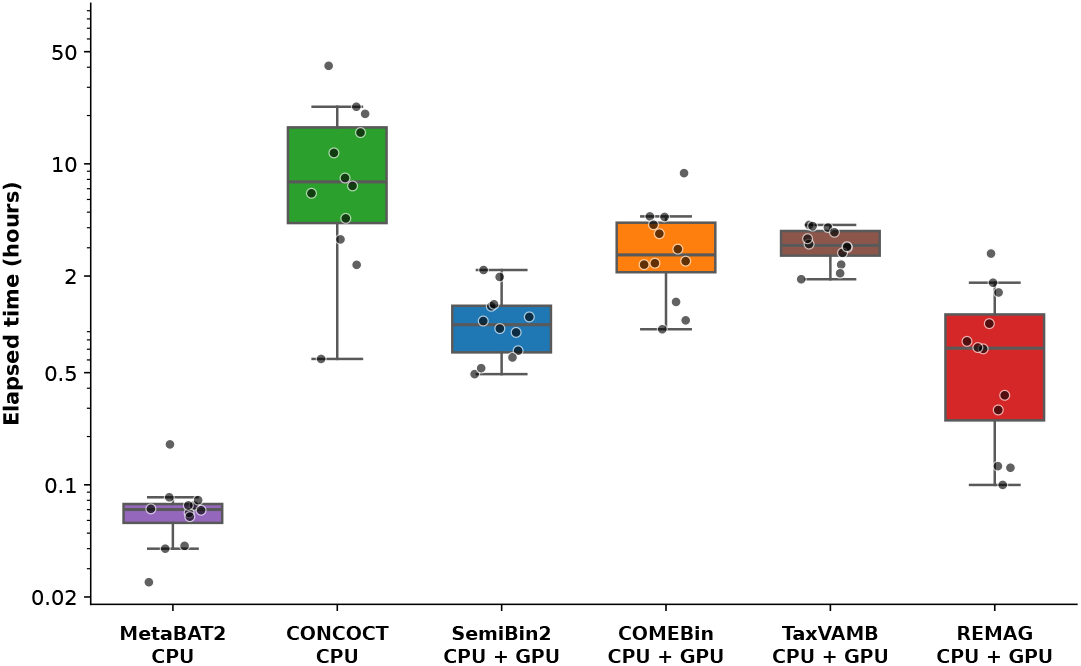
Runtime comparison across binning tools with GPU acceleration where supported. Elapsed execution time across 12 simulated co-assemblies from four datasets and three sequencing technologies. Each point represents one co-assembly. Boxes show the median and interquartile range, with whiskers extending to the most extreme observations within 1.5 times the interquartile range. All observations are shown on a logarithmic axis. Each run allocated 16 CPUs. SemiBin2, COMEBin, TaxVAMB and REMAG used GPU acceleration. MetaBAT2 and CONCOCT ran on CPUs. REMAG timings precede the filtering and encoder update.

**Figure S5.**
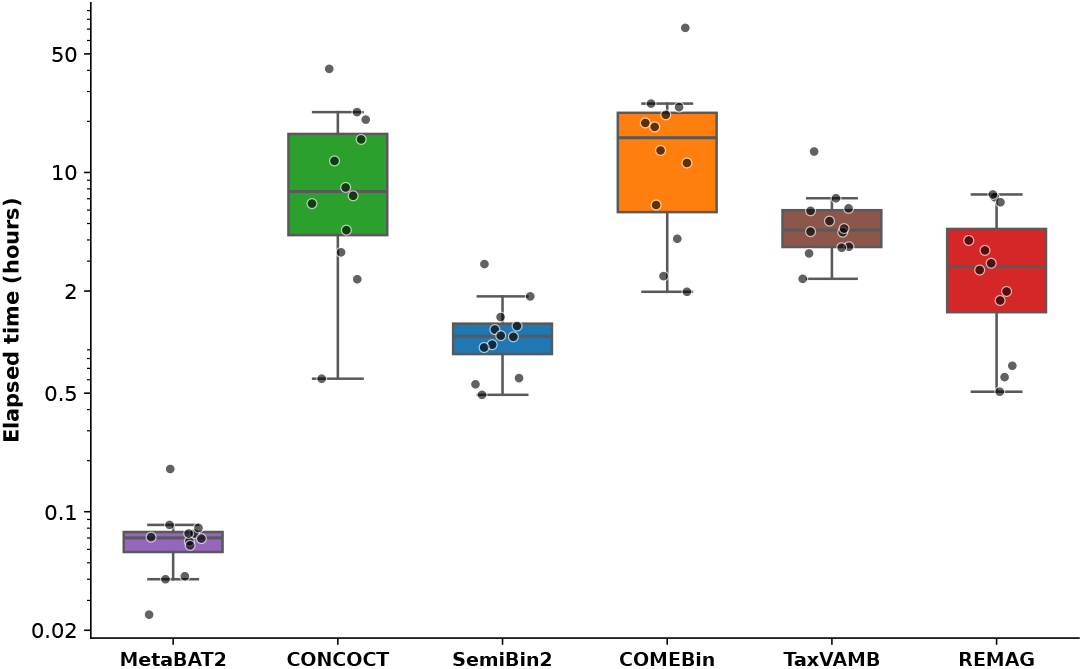
CPU-only resource use across binning tools. Elapsed time across 12 simulated co-assemblies, using 16 CPU cores per run. Points show individual co-assemblies and boxes show the median and interquartile range. REMAG timings precede the filtering and encoder update.

**Figure S6.**
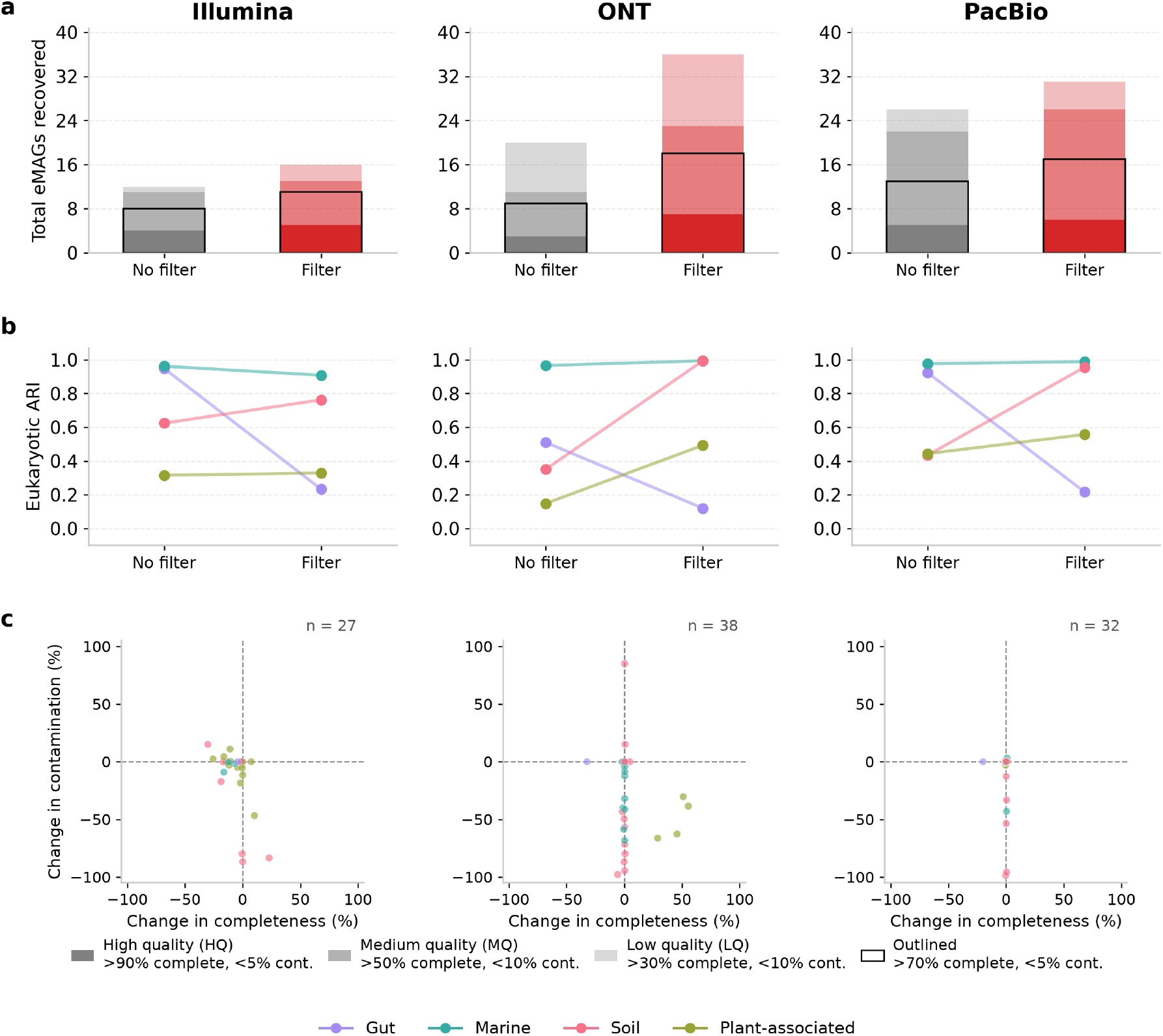
Ablation testing of eukaryotic contig filtering step. Comparison of REMAG with and without the eukaryotic contig filtering step on simulated gut, soil, marine and plant-associated datasets for the recovery of eukaryotic MAGs. Rows show (a) genome recovery, (b) eARI and (c) changes in completeness and contamination for matched reference genomes, calculated as filter minus no filter. Panel (c) includes matched references with *>* 30% completeness and *>* 200 kb of covered reference sequence in at least one condition.

**Figure S7.**
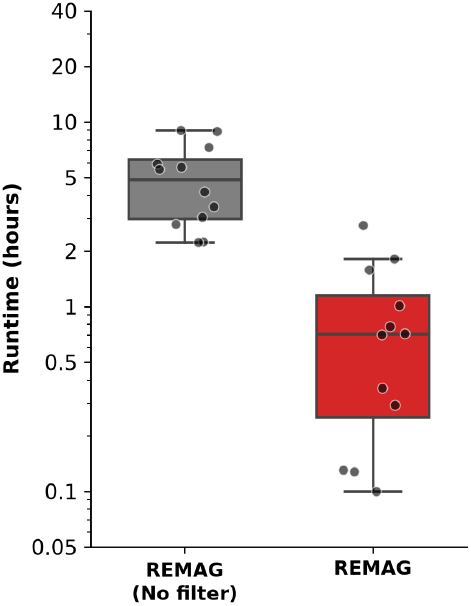
Impact of eukaryotic contig filtering on REMAG runtime. Comparison of GPU-assisted REMAG execution times with and without the eukaryotic contig filtering step across all simulated datasets.

**Figure S8.**
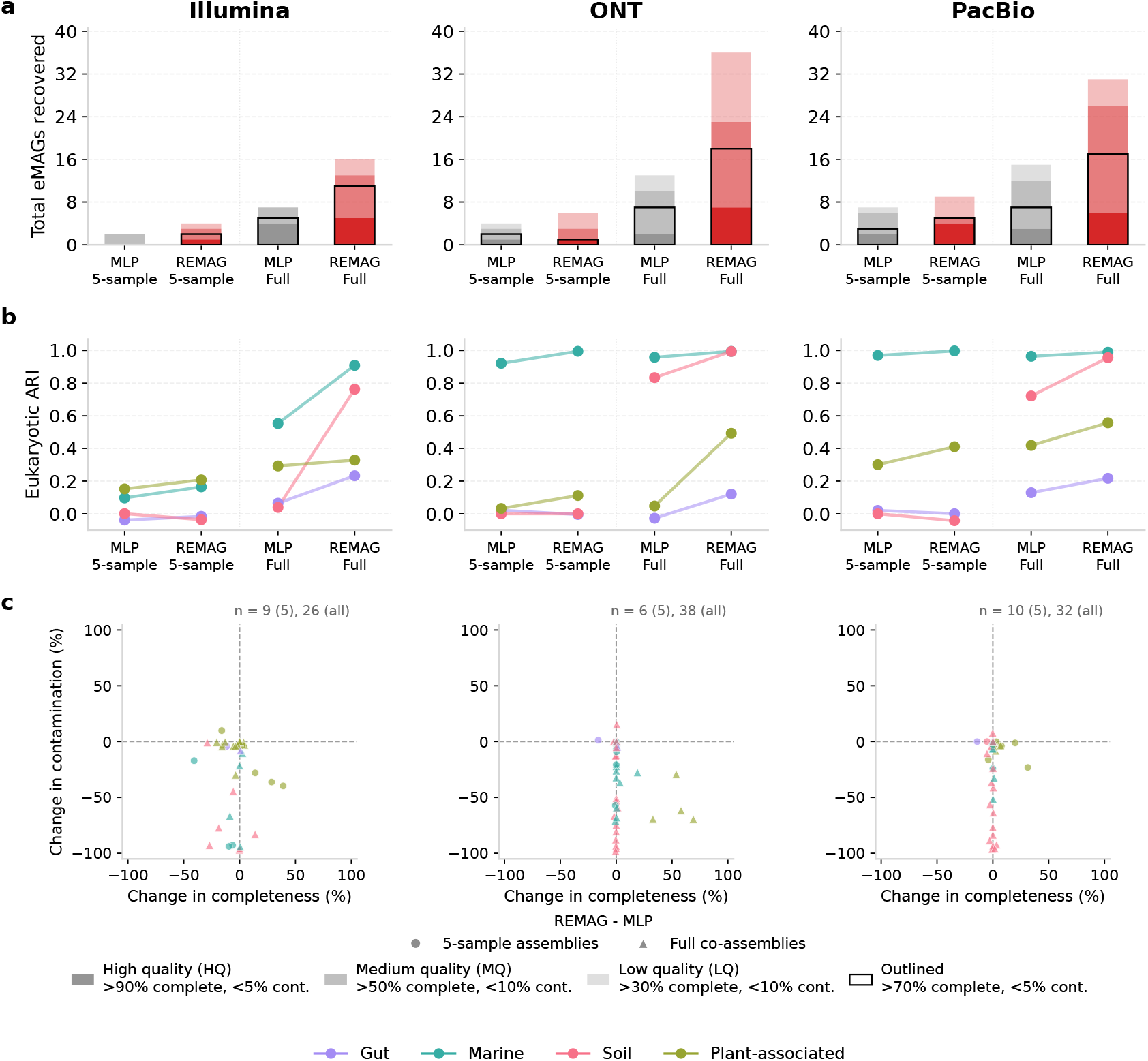
Ablation testing of the REMAG encoder. Comparison of REMAG and a simple multi-layer perceptron (MLP) encoder with 64 hidden units on full co-assemblies and independently constructed five-sample co-assemblies. Each encoder pair uses the same assembly and all its coverage profiles. Rows show (a) genome recovery, (b) eARI and (c) changes in completeness and contamination for matched reference genomes, calculated as REMAG minus the simple MLP. Circles and triangles in panel (c) distinguish five-sample and full co-assemblies, respectively. Panel (c) includes matched references with *>* 30% completeness and *>* 200 kb of covered reference sequence in at least one condition.

**Figure S9.**
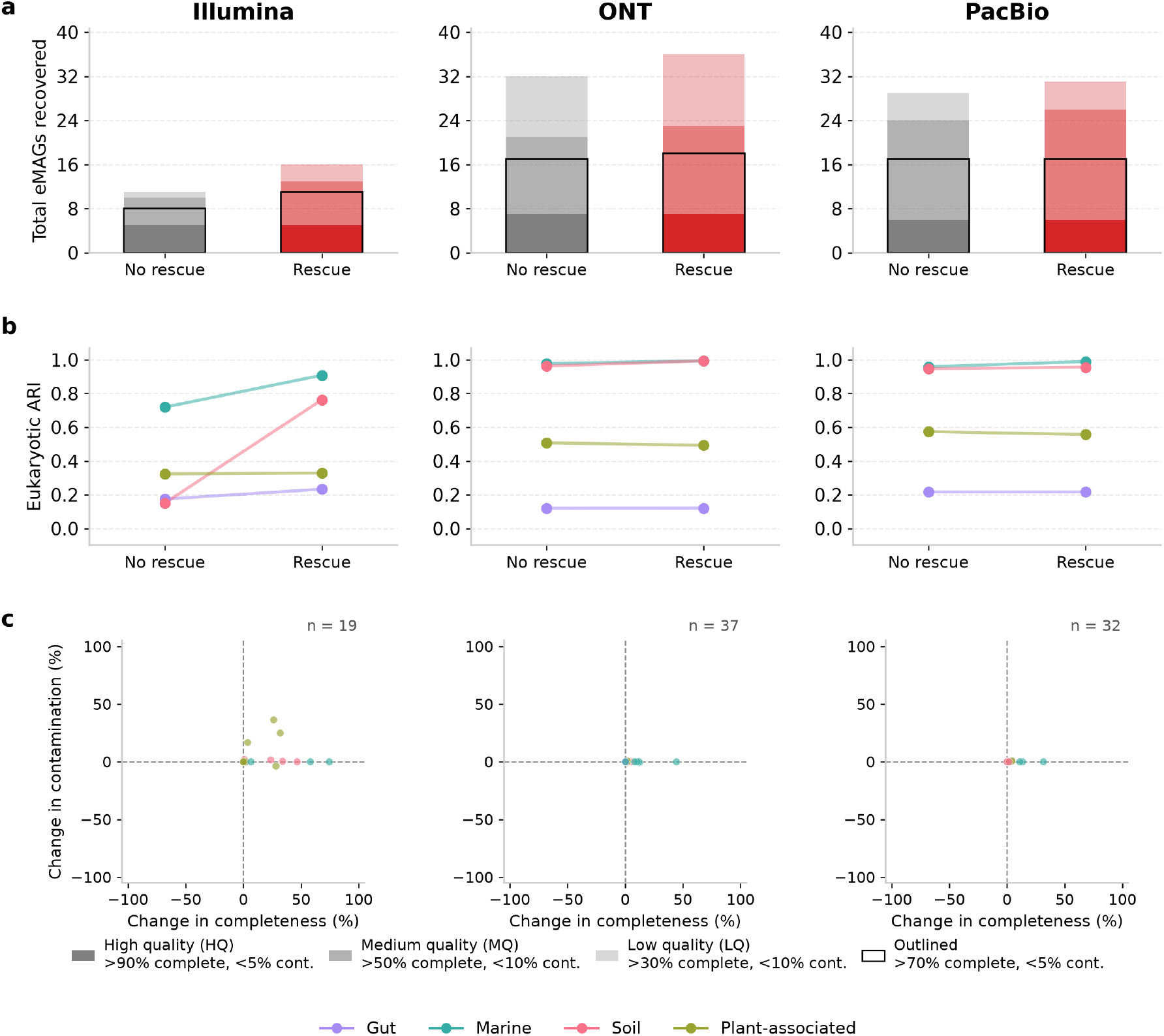
Ablation testing of unified rescue step. Comparison of REMAG performance with and without unified rescue, including both bin merging and contig recruitment, on simulated gut, soil, marine and plant-associated datasets for the recovery of eukaryotic MAGs. Rows show (a) genome recovery, (b) eARI and (c) changes in completeness and contamination for matched reference genomes, calculated as rescue minus no rescue. Panel (c) includes matched references with *>* 30% completeness and *>* 200 kb of covered reference sequence in at least one condition.

**Figure S10.**
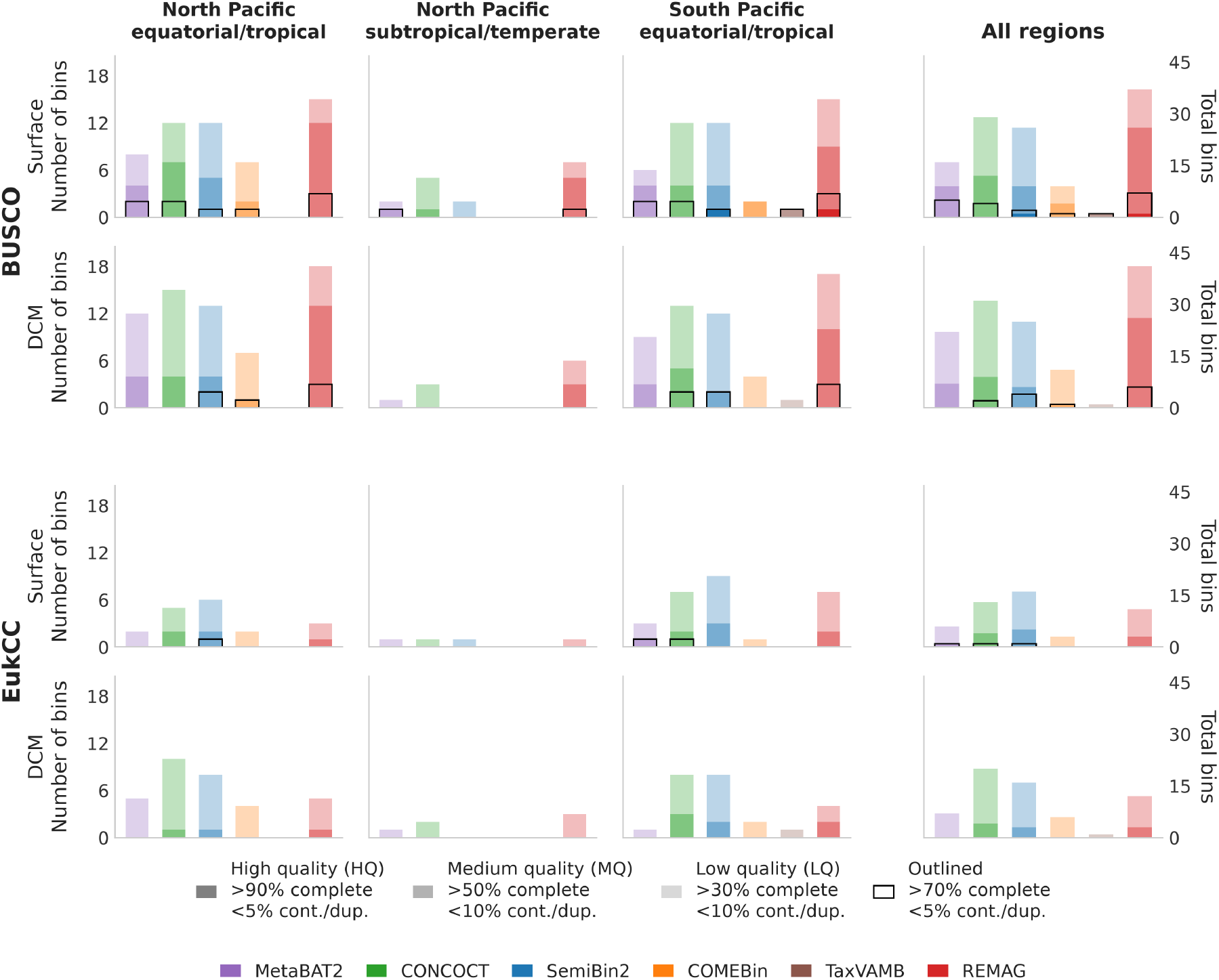
Eukaryotic MAG recovery performance on short-read *Tara* Oceans datasets. Comparison of MetaBAT2, CONCOCT, SemiBin2, COMEBin, TaxVAMB and REMAG across six *Tara* Oceans short-read coassemblies from three Pacific regions at surface and deep chlorophyll maximum (DCM). Bars show MAG quality tiers assessed by BUSCO (eukaryota_odb12.2) and EukCC. Right-hand panels show totals across regions, separately for each depth.

**Figure S11.**
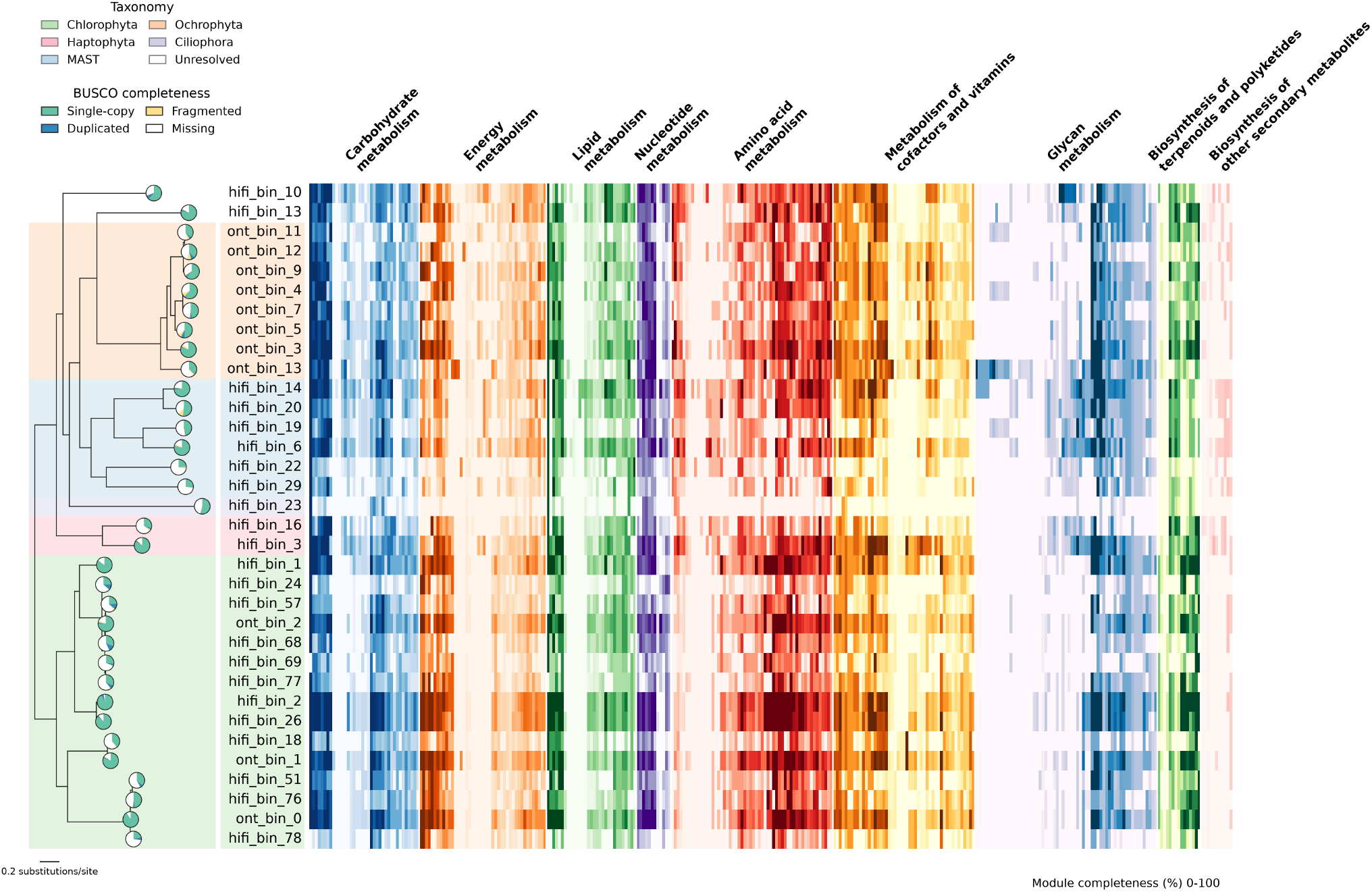
KEGG pathway completeness for plankton MAGs. Heatmap showing KEGG module completeness in the 34 plankton eMAGs in Figure 4. Modules are grouped by metabolic category. Genome BUSCO pies show the proportions of single-copy, duplicated, fragmented and missing markers.

## Tables

**Table S1.** Genome recovery and total elapsed runtime across simulated co-assemblies. Counts and runtimes are summed across 12 simulated co-assemblies. Ratios divide total HQ or HQ+MQ recovery by total runtime in hours. CPU + GPU indicates GPU-assisted execution.

| Method | Hardware | HQ | HQ+MQ | Total runtime<br>(hours) | HQ /<br>runtime hour | HQ+MQ /<br>runtime hour |
| --- | --- | --- | --- | --- | --- | --- |
| MetaBAT2 | CPU | 2 | 12 | 0.87 | 2.303 | 13.820 |
| CONCOCT | CPU | 12 | 32 | 144.27 | 0.083 | 0.222 |
| SemiBin2 | CPU | 5 | 31 | 14.31 | 0.350 | 2.167 |
| SemiBin2 | CPU + GPU | 5 | 31 | 13.10 | 0.382 | 2.366 |
| COMEBin | CPU | 7 | 28 | 220.80 | 0.032 | 0.127 |
| COMEBin | CPU + GPU | 7 | 28 | 39.52 | 0.177 | 0.708 |
| TaxVAMB | CPU | 12 | 32 | 64.14 | 0.187 | 0.499 |
| TaxVAMB | CPU + GPU | 12 | 32 | 37.79 | 0.318 | 0.847 |
| REMAG | CPU | 18 | 62 | 39.86 | 0.452 | 1.556 |
| REMAG | CPU + GPU | 18 | 62 | 10.38 | 1.734 | 5.971 |

**Table S2.** *Tara* Oceans sequencing runs used for short-read benchmarking. Paired-end Illumina sequencing runs used for six *Tara* Oceans co-assemblies from three Pacific regions at surface and deep chlorophyll maximum (DCM), for the 20–180 µm size fraction. All runs are available from the European Nucleotide Archive (ENA).

| Zone | Station | Sample accession | Run accession(s) |
| --- | --- | --- | --- |
| **North Pacific equatorial/tropical** |  |  |  |
| Surface | 106 | SAMEA2732548 | ERR1726525 |
| Surface | 109 | SAMEA2657008 | ERR1726788, ERR1726801 |
| Surface | 129 | SAMEA2732648 | ERR1726894 |
| Surface | 137 | SAMEA2732378 | ERR1726692 |
| Surface | 138 | SAMEA2732658 | ERR1726948, ERR1726967 |
| DCM | 106 | SAMEA2732775 | ERR1726830 |
| DCM | 109 | SAMEA2657103 | ERR1726563, ERR1726658 |
| DCM | 129 | SAMEA2732208 | ERR1726647 |
| DCM | 137 | SAMEA2732613 | ERR1726954 |
| DCM | 138 | SAMEA2732721 | ERR1726874 |
| **North Pacific subtropical/temperate** |  |  |  |
| Surface | 132 | SAMEA2732790 | ERR1726580, ERR1726872 |
| Surface | 133 | SAMEA2732917 | ERR4691819 |
| Surface | 135 | SAMEA2732980 | ERR1726750 |
| DCM | 132 | SAMEA2732910 | ERR1726769 |
| DCM | 133 | SAMEA2732861 | ERR4691814 |
| DCM | 135 | SAMEA2732990 | ERR1726641, ERR1726680 |
| **South Pacific equatorial/tropical** |  |  |  |
| Surface | 100 | SAMEA2657086 | ERR1726588, ERR1726639, ERR1726755 |
| Surface | 102 | SAMEA2657092 | ERR1726709 |
| Surface | 111 | SAMEA2657036 | ERR1726670 |
| Surface | 122 | SAMEA2657094 | ERR1726927, ERR1726932 |
| Surface | 127 | SAMEA2731923 | ERR4691798 |
| Surface | 128 | SAMEA2732035 | ERR1726577 |
| DCM | 100 | SAMEA2657077 | ERR1726655 |
| DCM | 102 | SAMEA2657087 | ERR1726659 |
| DCM | 111 | SAMEA2656980 | ERR1726690 |
| DCM | 122 | SAMEA2657064 | ERR1726606, ERR1726700 |
| DCM | 127 | SAMEA2731938 | ERR4691803 |
| DCM | 128 | SAMEA2731945 | ERR1726727 |

**Table S3.** MGnify study accessions used for the simulated datasets. The three biomes (marine, human gut, and soil) and their corresponding MGnify study and sample accessions used for generating taxonomic profiles and abundance distributions for the simulated datasets.

| Biome | MGNify Study | Sample Accessions |
| --- | --- | --- |
| Marine | MGYS00001869 | ERS1800270–ERS1800278 |
| Human Gut | MGYS00002207 | ERS2181189–ERS2181198, ERS2181200–ERS2181208 |
| Soil | MGYS00005779 | ERS6356179–ERS6356181, ERS6356183–ERS6356184, ERS6356186, ERS6356192, ERS6356197, ERS6356203, ERS6356207, ERS6356214, ERS6356217–ERS6356219, ERS6356222, ERS6356224–ERS6356225, ERS6356231, ERS6356237–ERS6356238 |

